# Querying functional drivers in primary murine naïve and memory T cells using RNP-mediated CRISPR-Cas9 Gene Deletion

**DOI:** 10.64898/2026.02.05.704062

**Authors:** Surojit Sarkar, Yevgeniy Yuzefpolskiy, Vandana Kalia

## Abstract

Cytotoxic CD8 T lymphocytes (CTL) are critical for the clearance of pathogenic cells via antigen-dependent cell lysis, thus providing protection against infectious diseases and cancers. CTLs represent one of the key targets of immunotherapies and vaccine design. While microarray and single-cell RNA sequencing of effector and memory CD8 T cells have identified several promising gene targets that may modulate CD8 T cell responses, their development is slowed down by the financial and time constraints related to the generation of germline knockout mice for further validation studies. Here we present a protocol for conducting efficient deletion of genes from activated effector, as well as resting naïve and memory primary murine CD8 T cells using RNP-based Crispr/Cas9 technology. This CRISPR modification of CD8 T cells was then adapted to study the effects of gene deletion in the context of acute memory differentiation as well as chronic exhaustion of CD8 T cells. To this end, we have titrated the necessary dose of antigen-specific CD8 T cells to study their differentiation in acute and chronic infections and confirmed the model by demonstrating rapid expansion of CRISPR mediated PD-1 ablated CD8 T cells in a chronic viral infection. Finally, by combining this methodology with a murine model of subcutaneous tumor challenge, this study provides a unique screening system for genes critical for mediating clearance of malignant tumor cells by CTLs. This study expands current technical capabilities for rapid evaluation of functional role of any candidate gene in CTL responses to infections and cancer through targeted gene deletion at any stage of CD8 T cell differentiation without the need for germline gene deletion mouse models.

## INTRODUCTION

T cells continuously circulate in the body, scanning for and protecting their host from invasive intracellular infections and cancers^1^. Understanding the signals that drive CD8 T cell differentiation into terminal effector and central memory fates^2–6^ and how chronic antigen stimulation drives T cell exhaustion^7–9^ has been critical for designing potent long-lasting vaccines and is driving the revolutionary field of personalized medicine with tumor-specific CAR T cell immunotherapy at the forefront of the industry. Through the advent of population and single cell sequencing of antigen-specific T cells, the field is moving towards a better understanding of the factors driving effector T cell differentiation^2,8,10^. To probe the functional relevance of these genes, the field has generated an extensive library of germline knockout mice. However, studying T cells isolated from knockout mice can have a myriad of compounding errors as CD8 T cells maturing in a knockout environment could be different due to extrinsic or intrinsic genetic effects. This challenge was partially addressed by the development of mice with LoxP flanked genes bred to cell-subset specific Cre-expressing mice^11–13^, which allow for a more eloquent approach to studying gene ablation. While these powerful techniques have laid the foundation of our understanding of T cell differentiation, the process is expensive and extremely time consuming.

The advent of retroviral delivery for gene knock-down has become a staple in immunology for assessing the function of candidate genes and how they directly control T cell differentiation *in vitro* and *in vivo*^14–16^. Furthermore, CRISPR Genome editing tool has revolutionized biological sciences by creating a simple method of targeted gene ablation^31–33^. In this study we have aimed to develop a rigorous workflow whereby one can query the role of relevant genes in naïve, effector, and memory CD8 T cells and combine these knockout modalities with existing *in vivo* models of T cell memory differentiation and exhaustion. The synthesis of these tools provides a dynamic range of CD8 T cell manipulation that has not limited thus far.

We have utilized the IDT CRIPSR-Cas9 platform combined with the Neon transfection system to knockout genes in naïve, primed effector, and memory CD8 T cells. We have demonstrated that the genes can be efficiently knocked out in effector CD8 T cells following optimal CD8 T cell priming, as well as in naïve and memory CD8 T cells following cytokine incubation. We further standardized the voltage conditions, cell-yield approximations, adoptive cell dose transfer, and time of transfer to rigorously study the antigen-specific CD8 T cell responses under conditions of acute memory differentiation, chronic viral exhaustion, as well as in the context of solid tumor clearance. To confirm the fidelity of the CRISPR/Cas9 methodology for targeted gene deletion in mature peripheral CD8 T cells in the context of chronic viral infection, we sought to replicate the findings from a published study demonstrating that germline PD-1^-/-^ CD8 T cells undergo unchecked expansion in chronic antigen milieu^17^. Our study compared WT, PD-1 CRISPR knockout and the same germline PD-1 knockouts that were used for the initial study. We found that CRIPSR/Cas9-mediated gene editing in CD8 T cells phenocopied the germline knockout cells. Finally, we were able to generate a high-throughput model that delineates genes critical in T cell mediated tumor clearance. As a proof-of-concept we utilized the MC38 solid tumor model and adoptively transferred WT and CRISPR PD-1 knockout tumor-specific CD8 T cells and observed a significant increase in tumor regression upon PD-1 ablation. This method offers a powerful platform for rapid and efficient deletion of candidate genes in primary murine CD8 T cells towards accelerating the preclinical pipeline of CAR-T cell engineering strategies that enhance solid tumor control.

## MATERIALS AND METHODS

### CRISPR-CAS9

Protocol for preparation of CRISPR-Cas9 reagents was followed closely to that provided by IDT Alt-R CRISPR-Cas9 System (www.idtdna.com/pages/support/guides-and-protocols). Briefly, crRNA and ATTO550-tracrRNA (keep protected from light) were resuspended in nuclease free IDTE buffer at 200μM and stored at -20°C until day of electroporation. crRNA and ATTO550 tracrRNA were melted together at a 1:1 mixture with a final concentration of both RNA at 44μM at 95°C for 5 min in a thermocycler. Products were then allowed to cool at room temperature for 15 minutes. Alt-R Cas9 enzyme was also stored at -20°C at 61 μM until day of experiment. Cas9 enzyme was diluted to 36μM in T buffer when using voltages greater than 1600 V and R buffer when using voltages less than 1600 V on the Neon System. Once the 44μM crRNA-tracrRNA mixture cooled to room temperature, it was combined 1:1 with the 36μM Cas9 solution, and incubated at room temperature for 15 minutes. The CAS9-tracr-crRNA were then combined with Electroporation enhancer at a 1:2 ratio (Complex:EPE). Electroporation enhancer was stored at 10.8μM at -20°C until the day of electroporation. Antigen-specific CD8 T cells were resuspended in R or T buffer, depending on necessary voltage, at 35x10^6^ cells/mL and combined at a 3:1 ratio of cells:Complex-EPE. Mixture was stored at room temperature prior to Neon electroporation.

### crRNA

crRNA was designed using the Broad Institute Genetic Perturbation Platform (portals.broadinstitute.org/gpp/public/analysis-tools/sgrna-design). The PD-1 crRNA used was as follows crRNA#1 CAATACAGGGATACCCACTA, #2 GACACACGGCGCAATGACAG, #3 CAGCTTGTCCAACTGGTCGG, #4 TTGGCTCAAACCATTACAGA, #5 GCACCCCAAGGCAAAAATCG. The CD25 crRNA used was GTGTCTGTATGACCCACCCG. The SATB1 crRNA used was as follows #1 GTACGTGCTGTTCACAATGG, #2 GCATCTGTCACATACGACAG. The CXCR3 crRNA used was as follows TCTGCGTGTACTGCAGCTAG. crRNA was resuspended at 200μM in nuclease free IDTE buffer and stored at -20°C.

### Neon Electroporation system

The Neon 10μL and 100μL kits (Invitrogen MPK1096) were used based on cell numbers. When fewer than 6x10^6^ cells, 10μL kit was used with 3 mL of room temperature E buffer in the Neon chamber. For greater than 6x10^6^ cells the 100μL kit with 3mL of room temperature E2 buffer in the Neon chamber was used. Naïve and memory CD8 T cells were resuspended in T buffer and electroporated using 2200V, 10ms, 3 pulses. CD8 T cells activated for 24-60hrs were resuspended in R buffer and electroporated using 1600V, 10ms, 3 pulses. Regardless of buffer, cells were resuspended at 35x10^6^ cells/mL before being mixed with their respective CRISPR-CAS9 reagents. Electroporated cells were pipetted into room temperature RPMI media (Thermo), made with 10% FBS (Gibco), 1% Penn-Strep/L-Glu (ThermoFisher), 0.4% BME (Sigma). Following electroporation cells were rested at 37°C, 5% CO_2_ for 30 minutes.

### CD8 T cell preparation

Direct *ex vivo* Neon Transfection of naïve and memory CD8 T cells results in large cell loss, and low transfection efficacy. To increase survival and frequency of transfection, first naïve and memory CD8 T cells were purified from spleen using a CD8 negative purification MojoSort kit (Biolegend). CD8 T cells were purified from naïve P14 mouse splenocytes, and memory CD8 T cells were obtained from B6 mouse splenocytes after day 60 post-LCMV_Arm_ infection. Following purification, naïve and memory CD8 T cells were plated in 6 well plates at 6x10^6^ cells/well in RPMI 10% FBS, 1% Penn-Strep/L-Glu, 0.4% BME and 10ng/mL mIL-7 (PeproTech)^18^. Cells were incubated at 37°C, 5% CO_2_ for 24 hours. As described previously, incubation with IL-7 increased T cell survival, as well as Neon transfection^18^.

To generate activated CD8 T cells, naïve CD8 T cells were purified using CD8 negative purification Mojo Kit from naïve P14 mouse spleens and activated *in vitro* with antigen presenting beads, or plates coated with αCD3/αCD28. Cells were plated at 2-3x10^6^ cells/well in 6 well plates in RPMI supplemented with 10% FBS, 1% Penn-Strep/L-Glu, 0.4% BME. Antigen-presenting beads were generated by incubating SA-Magnetic Beads (Sigma) with αCD3/αCD28-Biotin (Biolegend), or MHCI-GP33/αCD28-Biotin. Where indicated, αCD3(4μg/mL)/αCD28(4μg/mL) was also coated on F-Bottom plates for 2hrs prior to T cell activation at 37°C, 5% CO_2_, or O/N at 4°C. Prior to neon transfection, if APBs were used, magnetic beads were removed from T cells using Stem Cell/MojoSort magnet incubation for 2 min at room temperature.

Following Neon transfection antigen-specific CD8 T cells were transferred into warm RPMI supplemented with 10% FBS, 1% Penn-Strep/L-Glu, 0.4% BME. Cells were rested at 37°C, 5% CO_2_ for 20-30min as post-electroporation efficiency was checked. Naïve and effector CD8 T cells were then plated with APB or on αCD3/αCD28 coated plates for activation. Effector cells were restimulated in the presence of mIL-2 (10ng/mL). Memory cells were cultured with mIL-7/-15 (10ng/mL).

### Antibody Staining

All antibodies for FACS staining were purchased from Biolegend (San Diego, CA, USA) with the exception of SATB1 (Abcam). Cells were stained for surface markers in 1% PBS, 1%FBS, 0.2% Sodium Azide. Intracellular staining was performed using the BD intracellular staining kit. Intranuclear staining for SATB1 was performed using the FOXP3 staining kit (Invitrogen). Flow cytometry was performed on an LSRII Fortessa (BD Biosciences, San Jose, CA). FlowJo (TreeStar, Inc) was used to analyze and plot FACS data. For direct *ex vivo* analysis of T cells from spleens (SPL) and inguinal lymph nodes (iLN), SPL and iLN were first mechanically disturbed using frosted slides, and 0.83% ammonium chloride was used to remove RBC contamination in SPL. Following Lung perfusion with cold 1xdPBS, TILs and lungs (LNG) were isolated and cut into pieces. Tumors and lungs were processed in Collagenase A, and then TILs, LNG and liver (LVR) were pushed through a 70 micron cell strainer, followed by Percoll centrifugation to isolate the T cell containing buffy coat.

### Mice

C57BL/6J mice (Thy1.2/1.2) were purchased from Jackson Laboratory and maintained in house. Thy1.1+ P14 mice expressing the H-2D^b^GP33 TCR specific to the GP_33-41_ epitope of LCMV were fully backcrossed to C57BL/6 mice and were maintained in our animal colony. Mice were infected with 2x10^5^ PFU LCMV_Arm_ IP for acute challenge, and 2x10^6^ PFU LCMV_Cl-13_ IV for chronic challenge. To generate memory mice, 10^5^ antigen-specific P14 CD8 T cells were adoptively transferred into naïve B6 mice, which were subsequently infected IP with 2x10^5^ PFU LCMV_Arm_. To standardize dose of electroporated donors in chronic infections, Neon transduced effector CD8 T cells were adoptively transferred IV at varying doses into day 2 chronic LCMV_CL-13_ infection-matched recipient mice. To study effector T cells in the tumor setting, 10^6^ MC38 cells expressing GP_33-41_ peptide of LCMV were grown *in vitro* and transferred subcutaneously into the flank of naïve B6 mice. Once tumors were palpable at day 8 post-tumor challenge, Neon transfected CD8 T cells were adoptively transferred at 10^6^ cells per mouse IV. Tumors were measured every third day for three weeks. Tumor measurements were based on the formula [(larger length)x(smaller length)^2^]x(1/2) .

### Statistical analysis

Paired or unpaired Student’s t-test was used as indicated to evaluate differences between two group means. One-way ANOVA analysis with a Tukey post-test was used when comparing more than two groups. All statistical analyses were performed using Prism 5 and P values of statistical significance are depicted by asterisk per the Michelin guide scale: * (P ≤ 0.05), ** (P ≤ 0.01), *** (P ≤ 0.001) and (P > 0.05) was considered not significant (ns).

## RESULTS

### Protocol for screening genes critical for CD8 T cell function in naïve, effector or memory states

The goal of this protocol was to genetically modify antigen-specific CD8 T cells regardless of their state of differentiation using the Neon CRISPR platform. To achieve this, we first standardized proper preincubation conditions to make the T cells amenable to Neon-mediated CRISPR uptake for optimal gene deletion. The gene ablation was confirmed by following Neon transfected cells *in vitro* under conditions relevant to the cell state association of the gene being deleted (eg. genes upregulated upon activation were studied in T cells stimulated with antigen, while ablation of memory markers was confirmed with culture of cells in IL-7, IL-15 media to maintain quiescence). Following Neon CRISPR transfection, the transfected CD8 T cells will were transferred *in vivo* into the antigen milieu of choice to observe the effect of the gene knockout on CD8 T cell kinetics and function (Figure 1A).

**Figure 1.**
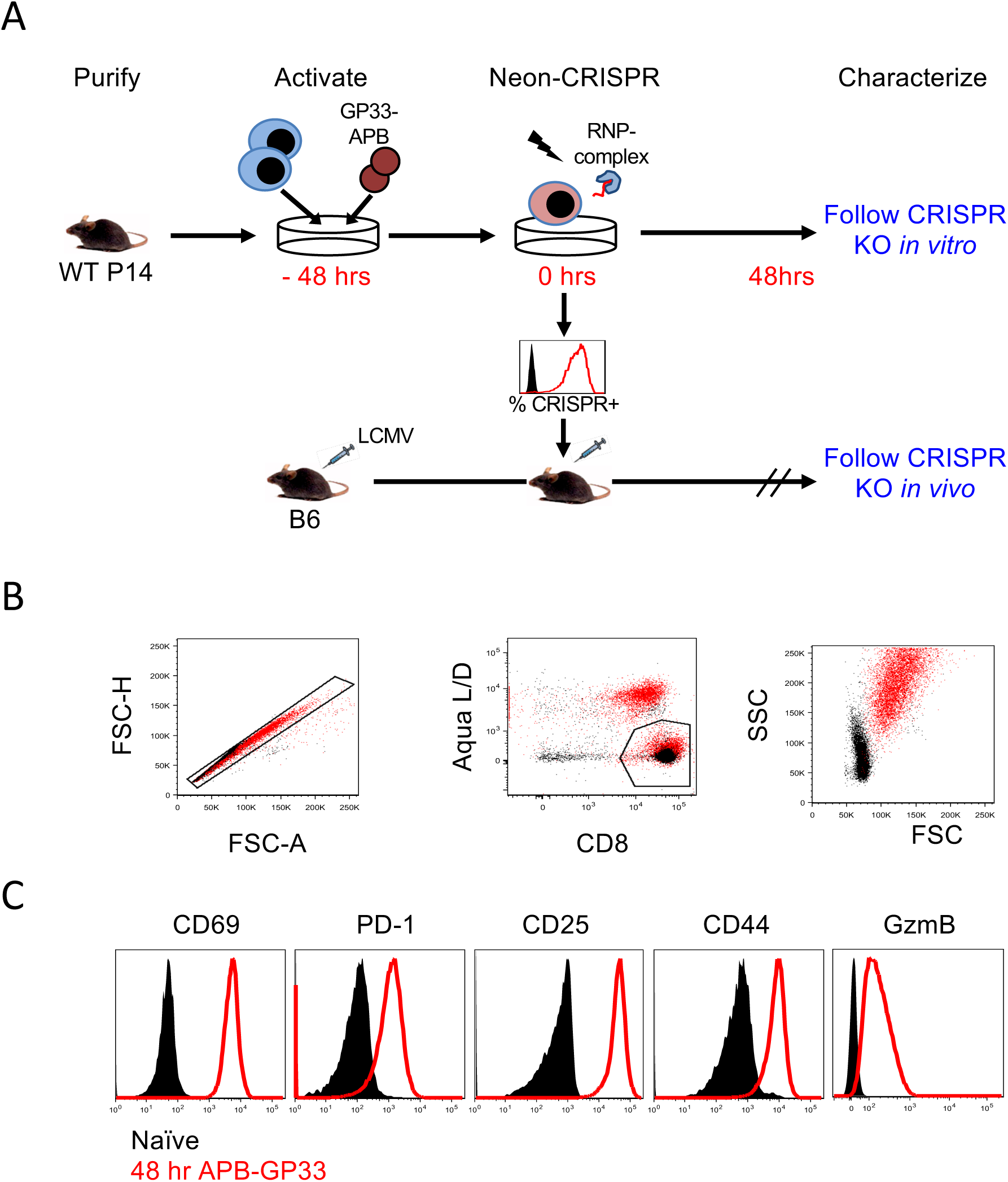
Optimal activation of antigen-specific CD8 T cells for CRISPR transfection. **A.** Experimental design. WT antigen-specific CD8 T cells are isolated from P14 mice and activated *in vitro* with antigen presenting beads coated with MHC-GP33 and αCD28. Cells are cultured for 36-48 hrs and magnetic beads are removed. Cells are resuspended in electroporation buffer together with CAS9 complex and electroporated using the Neon instrument. Cells are transferred into infection-matched mice and followed *in vitro* for knockout confirmation. B-C. Antigen-specific CD8 T cells are cultured *in vitro* with antigen presenting beads for 48 hrs. **B.** FACS plots show representative cell gating of naïve (black) and 48hr post-stimulation (red) antigen-specific CD8 T cells. **C.** Histograms depict activation markers of CD8 T cells 48 hrs post activation.

### Standardizing activation conditions for optimal uptake of Crispr/Cas9

We first standardized the *in vitro* activation conditions necessary for optimum uptake of CRISPR-Cas9 by CD8 T cells. As activated cells are fragile and amenable to death, it was important to develop a method of stimulation that required the least amount of handling. Thus, we magnetically purified CD8 T cells from naïve P14 spleens to prevent the need for purification at any later point in the protocol. To determine optimal CD8 T cells activation we applied two separate modalities – the first was stimulating with αCD3/αCD28 coated F-bottom plates, which provides a mono-interface for T cell activation and is widely used in the field. This plate coated stimulation for 48 hours provided efficient stimulation, as indicated by increased cell size (FSC), and upregulation of activation markers CD69, PD-1, CD25 and effector molecule Granzyme B (Figure S1C). However, to attempt a more physiologically relevant stimulation, we used SA-coated magnetic beads, which were bound with MHC-GP33 and αCD28 in equimolar concentrations. This provided a cell-cell approximate interface for activation, which is increasingly used in the field of CAR T cell immunotherapy expansion. Titration of these antigen-presenting beads (APB-GP33) demonstrated that a ratio of 1 APB for every two T cell yielded optimum activation in terms of activation marker expression, blast size as measured by FSC (Figure 1B, S1A), and yielded similar cell numbers compared to the other ratios tested (Figure S1B). Thus, for the future experiments where T cells needed to be activated, we utilized antigen-presenting beads at a 2:1 cell-bead ratio (Figure 1C).

### Optimizing voltage for CRISPR-CAS9 Uptake

The CRISPR-Neon protocol relies on the uptake of the CRISPR-CAS9 complex into cells via micropores opening upon the application of a voltage pulse through the cell solution. By utilizing the fluorescently labeled ATTO550-tracrRNA from IDT we were able to use flowcytometry to compare the uptake of CRISPR-CAS9 by GP33-specific TCR-transgenic P14 T cells. Standardization of different voltages revealed that activated effector CD8 T cells required a pulse of 1600V to uptake the majority of the CAS9 complex (Figure 2A). However, this uptake was paired with the loss of about 50% of the T cells directly after electroporation (Figure 2B). While this is a large cell loss it is important to have a high percent uptake as cell loss from sorting electroporated cells will be significantly greater (data not shown). Following electroporation, antigen-specific CD8 T cells were then maintained *in vitro* with antigen presenting beads for 48 hours, however, no additional toxicity was observed as the cells were still at about 50% survival compared to the 0V control (Figure 2C). Furthermore, we confirmed that the uptake of ATTO550 was not due to surface binding, as the fluorescence was maintained long after electroporation, and exhibited a gradual decrease in fluorescence overtime as the cells proliferated (Figure 2D). However, 48hrs following Neon transfection, the majority of the Cas9 complex was diluted from the system or broken down (Figure 2D).

**Figure 2.**
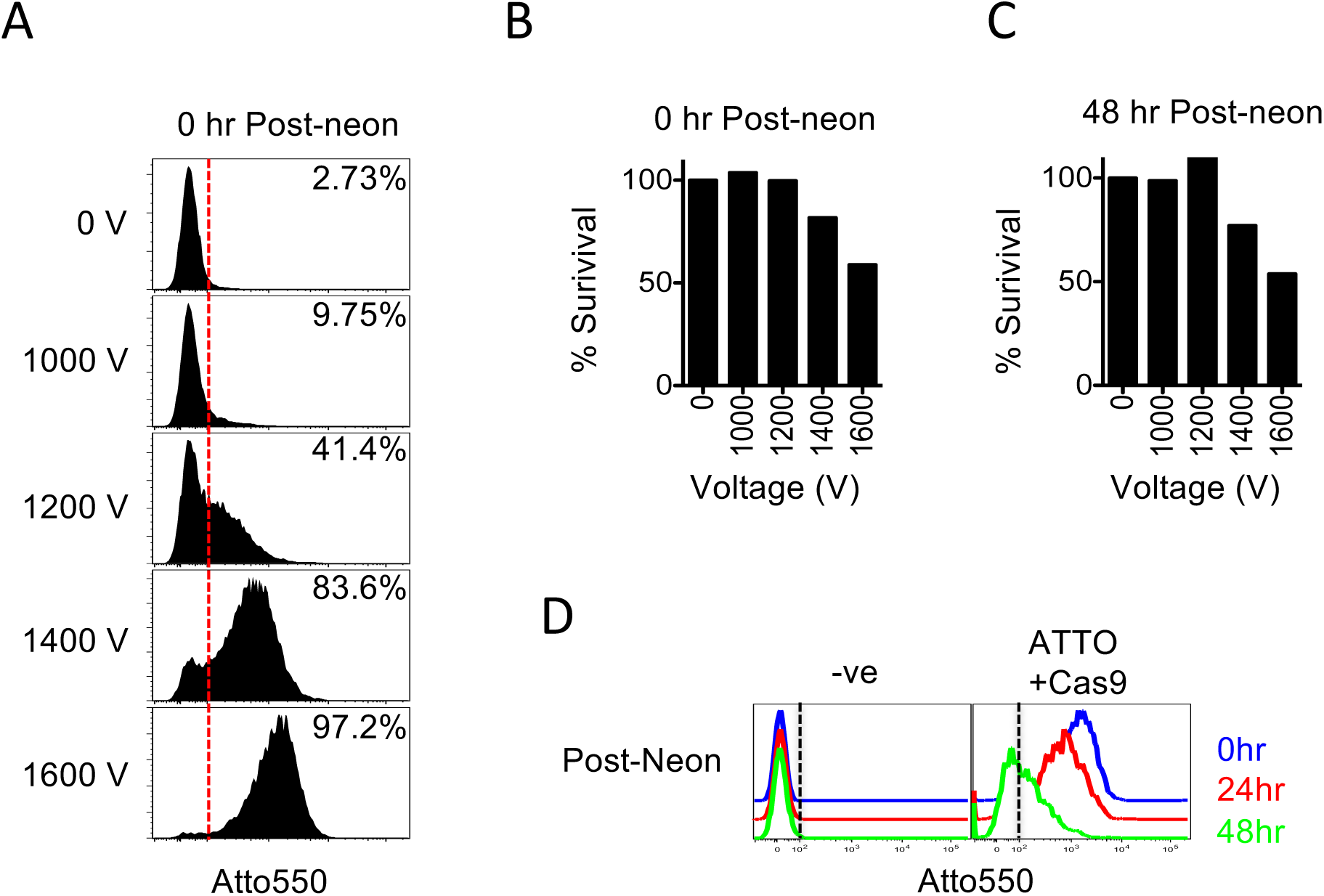
Activated CD8 T cells take up and retain CAS9 complex following electroporation. **A-B**. Antigen-specific CD8 T cells are electroporated following 48 hr stimulation. **A**. Histograms depict uptake CAS9 complex by fluorescence of tracrRNA-ATTO550. FACS plots depict survival of antigen-specific CD8 T cells following electroporation. **B**. Bar graph shows survival of antigen-specific CD8 T cells directly after electroporation. **C**. Following electroporation using 1600V antigen-specific CD8 T cells were cultured *in vitro* with APB-33+IL-2. Histograms depict ATTO550 at indicated times post electroporation.

In stark contrast to the activated T cells, direct *ex vivo* purified naive T cells were not as amenable to Neon transfection. Using the highest recommended voltage on the Neon resulted in only an 80% uptake of ATTO550 (Figure S2A). This electroporation resulted in a significant decline in CD8 T cells numbers directly after transfection (Figure S2B). The cells were then activated with GP33-APB and analyzed 48 hours later, where significant cell death was still observed (Figure S2C). The decreased uptake of Cas9 in naïve CD8 T cells was also paired with a dramatic decline in the observed CAS9 protein even just 24 hours after electroporation, with no observable CAS9 48 hours later (Figure S2D). This suggests that the majority of the ATTO550 expression observed was superficial coating on the cell surface of protein that was unable to enter the cell. To mitigate this issue of CAS9 uptake, we drew from previous studies using CD8 T cells, which showed that preincubation of resting CD8 T cells in IL-7 helped increase the fold-uptake of Cas9 product^18^. Thus, we utilized a similar preconditioning method on purified naïve CD8 T cells for 24 hours in IL-7. Notably, while we did not see a deviation from their resting phenotype (data not shown) we observed a dramatic increase in the uptake of CAS9 (Figure S2D).

### Choosing the appropriate crRNA target for protein knockout and validation

We next sought to determine if the increased ATTO550 fluorescence, which is our readout for CAS9 protein uptake, correlated with a functional CAS9 mediated gene deletion. To test the CAS9 mediated gene-knockout, we targeted a protein that is rapidly upregulated following T cell activation, PD-1. We confirmed that within the first 24-48 hours of activation, PD-1 was rapidly upregulated by effector CD8 T cells (Figure 1C, S1A, S1C). Furthermore, the wealth of data on PD-1 expression in chronic infections and tumors made it a perfect candidate for system validation^19,20^. Thus, we electroporated these activated T cells with ATTO-CAS9 and a PD-1 crRNA, and then cultured them for 2 days *in vitro* to asses PD-1 knockdown efficiency. We observed that PD-1 MFI and % positivity was drastically decreased with increasing voltage (Figure 3A) and directly correlated with the ATTO550 Cas9 uptake observed (Figure 2A). To determine which sgRNA to utilize, we engaged the Broad Institute Genetic Perturbation Platform and generated the top 5 best candidates. We demonstrated that all 5 candidates caused a significant decrease in PD-1 expression following Neon Transfection, leading to a decrease in PD-1 as early as 24 hours after Neon transfection, and nearly complete deletion by 48 hours following electroporation (Figure 3B). We confirmed that it was not a toxicity-mediated effect of Cas9-ATTO addition that caused the decrease in PD-1 expression, as both –ve (no neon) and ATTO+Cas9 (neon control) without PD-1 sgRNA had similar PD-1 expression (Figure 3C). Finally, to show that this method can target a wide variety of not only surface proteins but also transcription factors we targeted SATB1, shown to be relevant in antitumor function^21^, with the top two candidate sgRNAs. We observed that both targets showed pronounced decrease in SATB1 expression by 48 hours after electroporation compared to the electroporated control. (Figure 3D).

**Figure 3.**
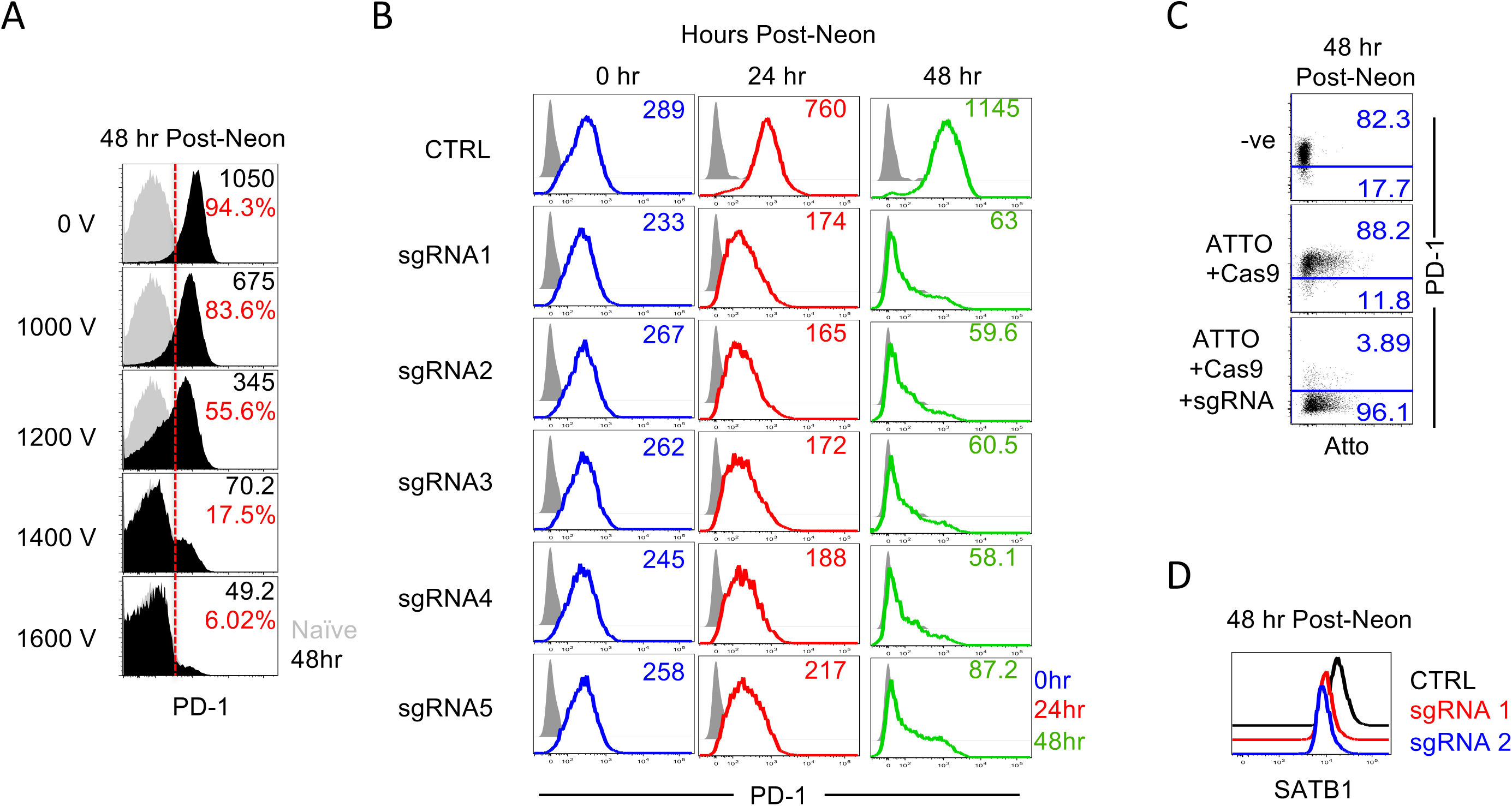
Incorporation of CRISPR following electroporation results in robust gene deletion. **A-D**. Antigen-specific CD8 T cells were stimulated for 48 hours using GP33-APB followed by Neon mediated electroporation with CAS9-crRNA. Cells were then transferred back *in vitro* and stimulated for 48 hours with APB-GP33. **A**. FACS plots depict PD-1 MFI following electroporation of CAS9-PD-1 crRNA (Black) compared to naive controls (grey) 48hours after Neon Transfection. **B**. FACS plots depict PD-1 MFI following electroporation using 1600V of top 5 PD-1 sgRNA candidates at given time post-electroporation. **C**. FACS plots show PD-1 expression as a function of CAS9 incorporation using 1600V for electroporation. **D.** FACS plots depict expression of SATB1 48hrs following electroporation of control (black) or two top candidate sgRNAs (Red and Blue) for SATB1.

To test the PD-1 knockout efficacy on naive CD8 T cells, we stimulated the T cells with antigen presenting beads for 48 hours following Neon transfection. As expected, naïve CD8 T cells electroporated with CAS9 and PD-1 crRNA only showed deletion in the highest voltage tested, however, even at this voltage we observed that the majority of cells still expressed PD-1 (Figure S3A). On the other hand, naïve T cells incubated in mIL-7 for 24 hours prior to electroporation had a dramatically decreased PD-1 expression by 48 hours after activation (Figure S3C). As CD25 also becomes drastically upregulated following CD8 T cell priming we used it as a knockout candidate to study the efficiency in CRISPR knock out of naïve IL-7 pretreated CD8 T cells. We observed that as with PD-1, although it still required a higher voltage of 2200 V, the majority of the cells were knocked out when pretreated with IL-7 (Figure S3B).

Finally, memory CD8 T cells are a hallmark of an optimum immune response, however their resting state makes it impossible to probe gene functions by gene editing, without a conditional knockout Cre-lox system. However, naïve and memory CD8 T cells share a large overlap in critical survival cytokines, with long-lasting memory cells upregulating their expression of IL-7Rα (CD127). Thus, we wanted to test if we could expand this naïve Neon CRISPR modality to knockout genes from memory CD8 T cells as well. To this end, we used spleens from LCMV memory mice to purify memory antigen-specific CD8 T cells utilizing CD44 as a marker of antigen experience^22^, and CXCR3 as a protein known to be upregulated in memory T cells^23^. We incubated the memory cells in IL-7 for 24 hours prior to transfection, and then performed Neon electroporation with Cas9 loaded with CXCR3 crRNA. We observed a significant decrease in CXCR3 expression in the CD44+ memory T cells (Figure S3D). We hypothesize that the remaining CXCR3 expression at day 3 post-Neon is due to the slow turnover of protein in resting memory cells as we observed a slow but steady decline from day 1 to day 3 post electroporation (Figure S3D).

Collectively, these studies demonstrate that this Neon CRIPSR protocol selectively knocks out surface and intracellular proteins from antigen-specific CD8 T cells prior to infection, during effector differentiation, and in resting memory cells. This provides a powerful toolset to study CD8 T cells at all stages of their development during immune responses.

### Comparison of established knockout model with CRISPR mediated knockout

Having standardized an efficient CRISPR-CAS knockout system, we next sought to confirm that the CRISPR-mediated gene deletion and protein loss led to physiologically relevant outcomes compared to established germline gene deletion models. Recent studies utilizing PD-1^-/-^ P14 T cells in chronic infections have shown that in the absence of PD-1, CD8 T cells undergo a vigorous expansion compared to their WT counterparts^17^. Thus, to understand how well our CRISPR model aligns with the germline gene deletion model, we first needed to standardize how many CRISPR modified T cells must be transferred into an infected mouse to obtain a measurable T cell response without causing supraphysiological levels of donors. To standardize the appropriate T cell dose, we cotransferred *in vitro* activated non-electroporated (control) cells with varying amounts of electroporated but non-CRISPR containing (Neon) T cells into infection-matched LCMV_CL-13_ infected mice. Mice were followed to the peak of effector expansion (day 8) and spleens were analyzed for donor P14 CD8 T cell responses compared to endogenous GP33-specific CD8 T cell responses. We observed that an adoptive transfer of 2x10^5^, *in vitro* 48hr activated, antigen-specific CD8 T cells into day 2 infected mice was able to match the endogenous frequency of GP33-specific CD8 T cells (Figure S4A-B). However, (as we showed previously) the electroporation process caused a significant amount of cell death, which was replicated *in vivo* as well, and it required 400-500k Neon antigen-specific CD8 T cells to match the number of control donors (Figure S4A-B). The transfer of donor cells did not impact the exhaustion phenotype of the endogenous or donor CD8 T cells as evidenced by expression of high levels of PD-1 at day 8 post-infection, typical of a chronic viral infection (Figure S4C).

Utilizing these standardizations, we performed PD-1 CRISPR knockout on WT (Thy1.1/1.1) *in vitro* activated CD8 T cells, and cotransferred them with WT electroporated control (Thy1.1/1.2) cells into day 2 LCMV_CL-13_ infected B6 mice (Figure 4A). A parallel experimental arm was setup to recapitulate the previous findings; containing *in vitro* activated germline PD-1^-/-^ P14 T cells (Thy1.1/1.1) cotransferred with WT electroporated control (Thy1.1/1.2) cells into day 2 LCMV_CL-13_ infected B6 mice. Utilizing the Neon conditions established in Figure 2, we confirmed that all cells effectively took up the ATTO550-Cas9 protein (Figure 4B). Furthermore, we demonstrated that prior to CRISPR mediated deletion of PD-1 both WT groups expressed similar levels, and only the somatic PD-1^-/-^ T cells expressed basal levels of PD-1 (Figure 4B). We confirmed the deletion of PD-1 *in vitro* at 48hrs post-Neon transfection and observed that the knocked-out cells had expression levels similar to PD-1^-/-^ CD8 T cells that were electroporated with just ATTO-Cas9 (Figure 4B). WT and PD-1 knockout CD8 T cells had similar starting ratios (Figure 4D), and were still comparable by day 8 post-infection. Following day 8 post-LCMV infection we observed a dramatic expansion in PD-1 knockout CD8 T cells in both the CRISPR knockout, and somatic knockout CD8 T cells, resulting in increased number of cells detected in PBMC of infected animals (Figure 4D). We confirmed that PD-1 was deleted even *in vivo,* with PD-1 CRISPR knockout cells showing baseline levels of PD-1 in spleen, but still expressing effector molecule Granzyme B. These data demonstrate that PD-1 CRISPR did not impact effector differentiation significantly (Figure 4E). We next compared CD8 T cell expansion in secondary lymphoid tissues as well as peripheral sites of infection, and observed a significant expansion of PD-1 CRISPR knockout donor cells in all tissues (Figure 4F). This recapitulates the previously published role of PD-1 mediated suppression of exhausted CD8 T cells^17^.

**Figure 4.**
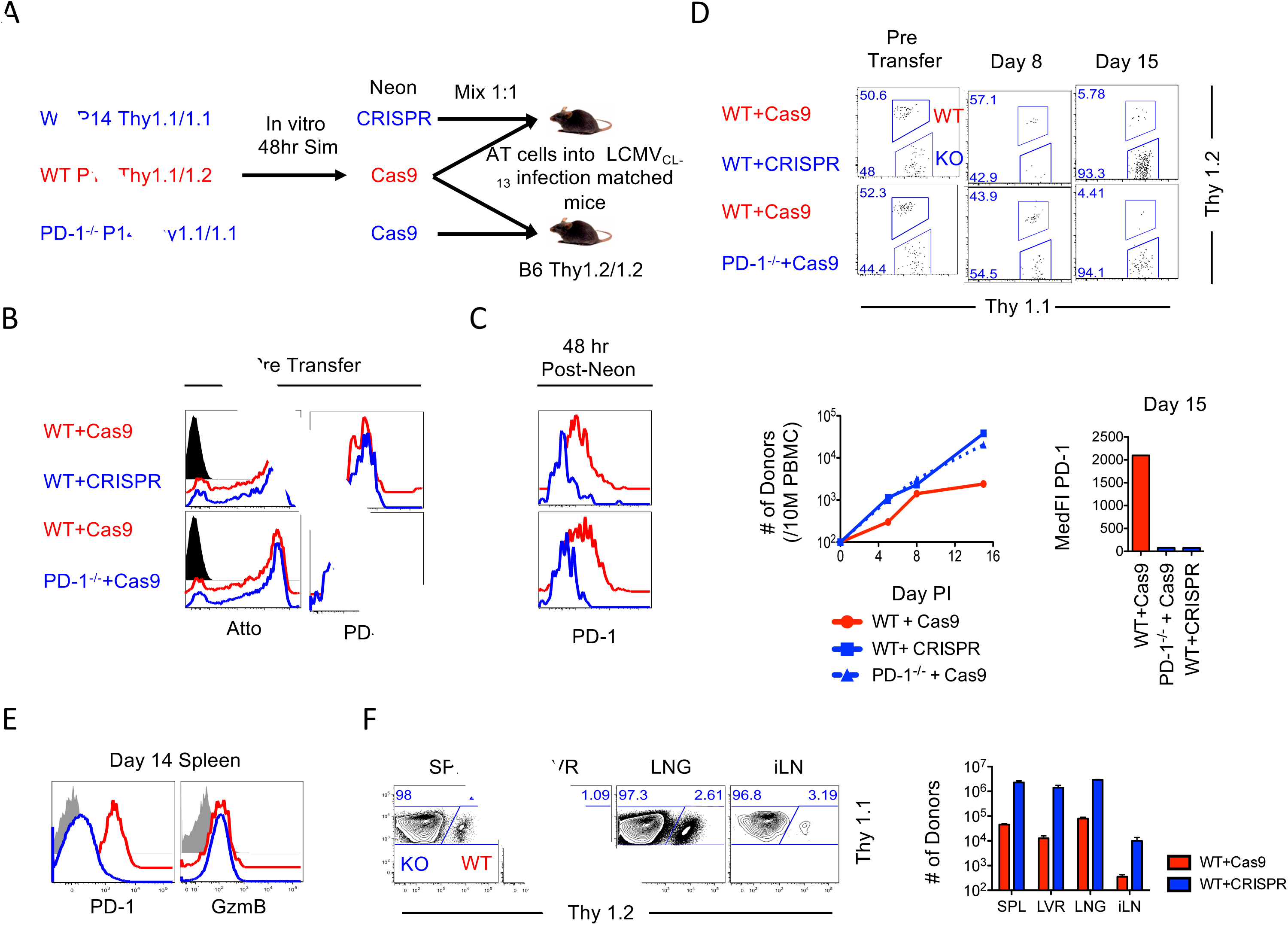
CAS9 gene deletion of PD-1 phenocopies germline PD-1^-/-^ CD8 T cells during chronic infection. **A**. WT or PD-1^-/-^ antigen-specific CD8 T cells were activated using APB-GP33 for 48hrs. Antigen-specific CD8 T cells were electroporated with CAS9 (control) or CAS9-PD-1 (CRISPR PD-1 knockout). WT control and WT CRISPR cells or WT control and PD-1KO control cells were cotransferred into infection matched LCMVCl-13 B6 mice. **B**. Histograms depict electroporation success and PD-1 expression prior to transfer into infection matched mice. **C**. Following electroporation cells that were not transferred *in vivo*, were cultured *in vitro* with APB-GP33+IL-2. Histograms depict expression of PD-1 48 hrs after electroporation. **D.** FACS plot shows relative ratio of WT Cas9 and WT CRISPR cells or WT Cas9 and PD-1KO Cas9 cells prior to transfer into infection matched B6 mice. FACS plots show relative ratio of donors at day 8 and 15 post infection. Line graph shows number of respective donor cells in blood of infected animals. Bar graph shows MFI of PD-1 in respective donor CD8 T cells at day 15 post-infection. **E**. Infected mice were analyzed at day 14 post-infection and spleen, iLN, LVR, and LNG were collected. Histograms show marker expression of PD-1 and GzmB at day 14 post-infection gated on donor WT-CAS9 (Red) and WT-CRISPR (Blue) cells, and naïve endogenous CD8 T cells (gray). **F.** FACS plots depict relative ratio of WT-Cas9 (WT) or WT-CRISPR (KO) in respective tissues. Bar graphs enumerate donor CD8 T cells numbers.

### Study of CRISPR transfection protocol on memory differentiation

We next wanted to confirm that we could also recapitulate normal memory differentiation following Neon electroporation. Thus, we electroporated *in vitro* activated CD8 T cells and adoptively transferred them into acute LCMV_Arm_ infection-matched mice, where memory cells develop after effective antigen clearance. At all doses studied, donor cells underwent vigorous effector expansion, differentiating into memory precursor and short-lived effector cells (MPEC and SLEC) at ratios consistent with previously published data (Figure S4D)^2,6^. We then followed these cells into the memory phase. Long-lived memory CD8 T cells were observed at day 30 post-infection (Figure S4E). Furthermore, we noted that Neon electroporated naïve CD8 T cells preconditioned with IL-7 for 24 hours, could be transferred at numbers identical to their direct *ex vivo* naïve transferred counterparts, resulting in similar effector expansion and memory differentiation kinetics (data not shown). These data establish robust application of the CRISPR-CAS9 knockout methodology to study gene function during effector CD8 T cell expansion and memory differentiation.

### Utilizing CRISPR/Cas9 gene editing to predict functional relevance in tumor environment

Being able to recapitulate CD8 T cells dynamics under acute and chronic conditions following CRISPR editing, led us to evaluate whether this methodology could be utilized to study T cell dynamics in the localized exhausted environment of solid tumors. Between the advent of checkpoint blockade therapy, and the later development of CAR T cell immunotherapy, there is a growing impetus for discovering targets critical for driving effector CD8 T cell functions and preventing tumor-mediated exhaustion for effective clearance of malignant tissue. PD-1 checkpoint blockade therapy has been shown to drive powerful T cell functional reinvigoration in both chronic viral infections and solid tumors^19,24^. MC38 is one such tumor that is highly amenable to PD-1/PDL1 checkpoint blockade therapy^24^. Thus, to test protein function in a tumor environment, we started by knocking out the lead candidate, PD-1, from activated effector cells and comparing how ablation of PD-1 inhibitory signals in antigen-specific T cells impacts tumor control. We generated an MC38 cell line expressing LCMV GP33-41 peptide, established the tumor growth kinetics *in vivo* (Figure 5A). We then compared the impact of transferring activated effector CD8 T cells that were electroporated with ATTO (control) or ATTO with PD-1-CAS9 (knockout). We separately transferred control or knockout cells into mice with established tumor burden, and compared their relative ability to drive tumor recession (Figure S5A). We observed that donor ATTO CD8 T cells were able to stabilize the tumor and prevent further tumor growth, thus supporting the benefit of transferring activated effector CD8 T cells (Figure 5A). However, the adoptive transfer of PD-1 CRISPR knockout CD8 T cells was able to suppress tumor growth and mediate potent tumor recession (Figure 5A). We observed that these effects were not due to differential survival between WT and CRISPR KO CD8 T cells during transfer, as both populations were able to maintain similar frequencies in PBMC (Figure 5B). These peripheral cells had similar resting levels of activation markers and cell size in the spleen at day 24 post-tumor implant (Figure S5B). This resting phenotype in the donors was consistent with their high polyfunctionality, consistent with the thesis that CD8 T cells in the periphery are not being exposed to the tumor antigen (Figure S5D). However, when comparing donor cells isolated from tumors, we observed that CRISPR knockout CD8 T cells were unable to upregulate PD-1 in the tumor environment, albeit, they still expressed similar levels of activation marker CD69, exhaustion marker TIM3, and had similar size as WT cells (Figure S5B). When comparing the function of PD-1 KO T cells, we observed that in the absence of PD-1 signaling, donor TILs had slightly higher polyfunctionality with higher levels of IFN-γ, TNF-α and IL-2 expression following direct *ex vivo* stimulation (Figure 5D). This is consistent with the known effects of PDL1 blockade in chronic infections. However, despite differences in CD8 T cell function and tumor clearance, we did not observe differences in total cell number accumulation in spleen or TIL, nor an increase in the density of donor PD-1KO CD8 T cells compared to WT CD8 T cells in the tumors (Figure 5C, S5C). This suggests that while knocking out PD-1 increases their effectiveness in tumor clearance, it does not prevent the exhaustion of CD8 T cells. Instead, loss of PD-1 merely delays exhaustion, as has been suggested previously during chronic infection^17^. Together, these data demonstrate a useful model of CRISPR Cas9 gene editing in a tumor setting, which may be effectively leveraged to screen for gene functions in T cells in a solid tumor setting, with multiple readouts including tumor cell growth, phenotypic analysis of donor cells, and functional cytokine profile of exhausted TILs and peripheral circulating T cells.

**Figure 5.**
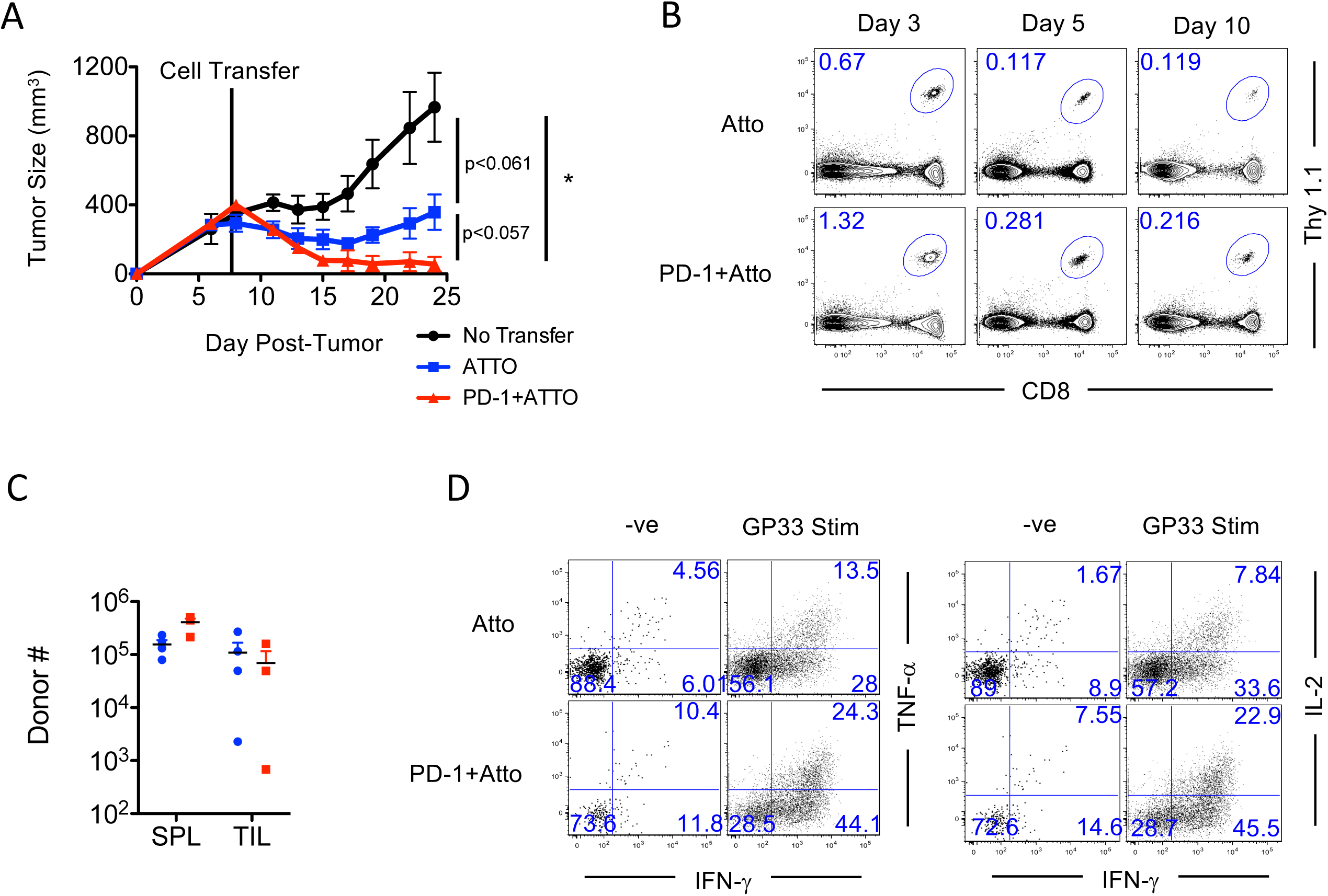
CRISPR model for determining genes critical in mediating CD8 T cell antitumor function. Naïve CD8 T cells were purified from spleens and stimulated for 48 hours using GP33-APB. Cells were electroporated with ATTO alone (control) or CAS9-PD-1-ATTO (PD-1 knockout) and transferred into tumor-burdened mice. **A**. Line graph depicts tumor size in mice, which received no donors (No transfer), WT effectors (ATTO), or PD-1 KO effectors (PD-1+ATTO). **B.** FACS plots show frequency of donor CD8 T cells in blood of recipient mice at given times post T cell transfer. **C**. Mice were carries to day 24 post-infection and tumors and spleens were collected. Graph depicts number of donor cells in spleen and TIL. D. TILs were stimulated for 5 hours in the presence of GP33 and BFA to measure cytokine functionality. FACS plots are gated on donor CD8 T cells in TILs.

## DISCUSSION

This study lays out the groundwork for studying effects of gene knockout in antigen-specific CD8 T cells during distinct phases of differentiation, without significantly perturbing them from their naïve, effector or memory states. We were able to demonstrate that this model works during acute infections, with CRISPR control cells undergoing standard effector expansion and memory differentiation. Furthermore, using PD-1 CRISPR KO CD8 T cells we were able to recapitulate the exaggerated T cell expansion during chronic infection, phenocopying the corresponding germline PD-1^-/-^ CD8 T cell phenotype. To increase the utility of this process we also titrated the number of donor CD8 T cells necessary to match endogenous frequencies following chronic infection, as well as the donors transferred during acute infection to match specific T cell benchmarks for day 8 effector expansion and memory formation.

The murine model of solid tumors is rich with a wide array of tumor-specific T cell mouse lines, as well as a multitude of different types of tumors such as glioma, melanoma, colorectal, and breast cancer lines, to name a few. In each scenario antigen-specific CD8 T cells respond uniquely and have different requirements to undergo sufficient rescue and tumor growth suppression. Using this Neon CRISPR protocol, we have demonstrated that we can target the genetic ablation of any target protein, and evaluate its effect on CD8 T cell mediated tumor growth kinetics. The difference between untransformed tumor growth, and the delayed tumor growth observed upon transfer of activated CD8 T cells demonstrates that there is flexibility in this model to further evaluate negative and positive gene regulators of antitumor responses. In fact, this study is amongst the first to study the effect of transferring PD-1^-/-^ effector CD8 T cells into MC38 tumor bearing mice, and demonstrating increased clearance that these cells were able to impose. Furthermore, we observed that despite lacking the PD-1 inhibitory receptor, these cells were still significantly exhausted compared to their peripheral circulating T cell counterparts in the context of solid tumors. These data are consistent with previous reports in the chronic LCMV infection model^17^. This experimental model system creates an exciting platform for deconvoluting signals necessary for understanding CD8 T cell immunosuppression in chronic environments.

Recent studies have demonstrated that Tox1/2-NR4A signaling axis is a critical mediator in the differentiation of exhausted CD8 T cells, and that a knockout of Tox1/2 could be beneficial in the clearance of tumor specific T cells^25,26^. Utilizing the Neon CRIPSR methodology of gene editing in primary murine T cells laid out here could help test multiple potential targets in parallel to compare localization, effector function, and polyfunctionality following genetic perturbation of the antigen-specific CD8 T cells in various stages of differentiation (naïve, effector memory). We believe the tools we have standardized here, along with similar recent advances in human T cells as well^27–30^, will be broadly applicable in T cell immunology, and can accelerate the field in determining genes critical for potent CD8 T cell responses to acute and chronic viral infections and well as solid tumors.

## ACKNOWLEDGMENTS

The authors would like to thank Dr. Hozefa Bandukwala for scientific input. This work was supported by NIH, NIAID (5R01AI132819 to SS) and seed funds from Seattle Children’s Research Institute (to VK and SS). Author Contribution: YY carried out experiments, analyzed data, prepared figures and wrote manuscript. SS and VK conceptualized the project, designed the experiments, carried out experiments, supervised the work, analyzed data, interpreted the results and wrote the manuscript. Conflict-of-interest disclosure: The authors declare no competing financial interests.

## SUPPLEMENTARY FIGURES

**Supplementary Figure 1.**
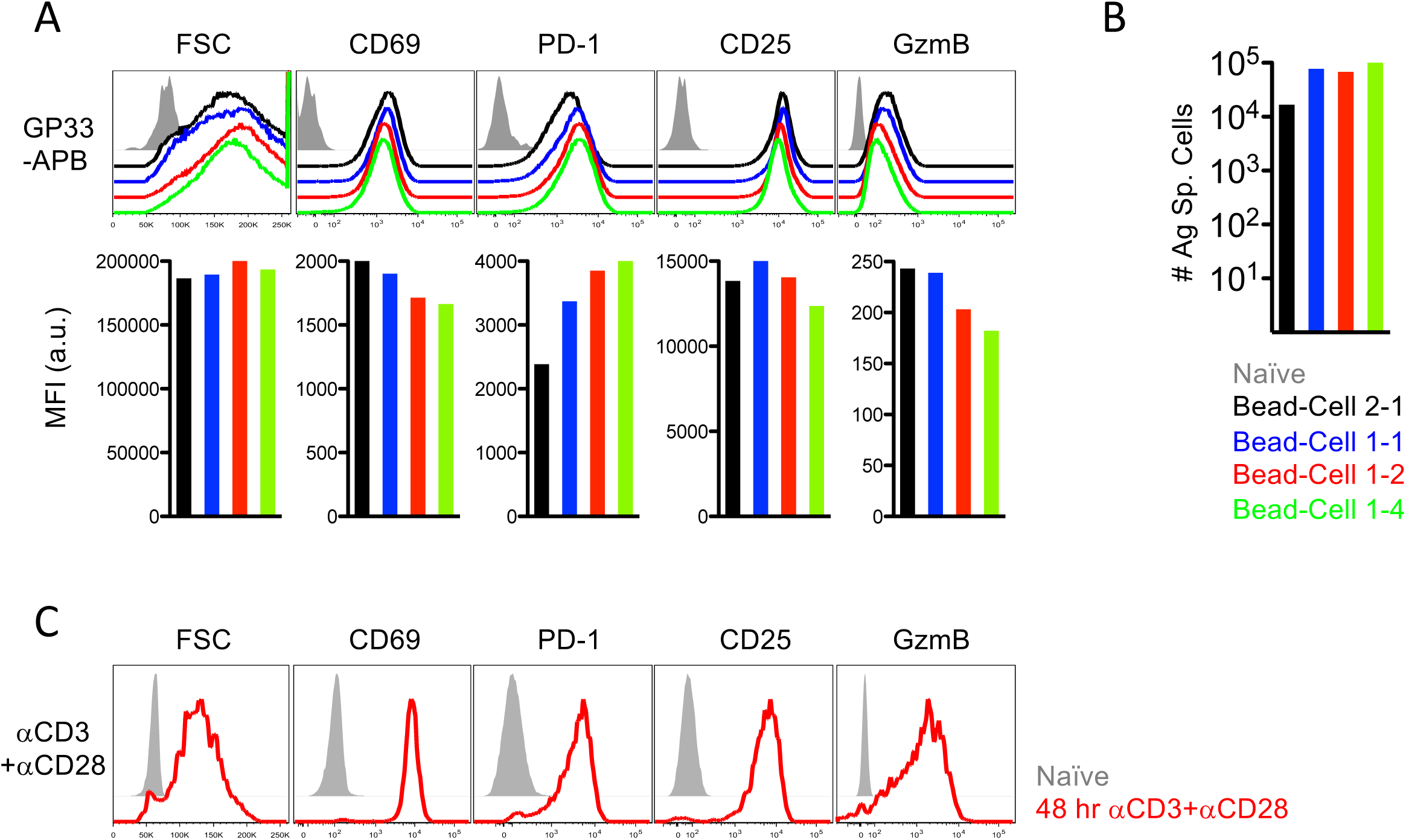
Titration of antigen presenting beads for in vitro priming of antigen-specific CD8 T cells. A. Antigen-specific CD8 T cells activated with different APB-CD8 T cell ratios. FACS plots depict markers after 48hrs in vitro stimulation. Bar graphs show MFI of respective markers. B. Bar graph shows number of antigen-specific CD8 T cells following 48 hr stimulation. C. Histograms depict activation markers following 48 hours of activation (red histogram) using anti-CD3/anti-CD28 coated plates. Gray histograms show expression in naïve T cells.

**Supplementary Figure 2.**
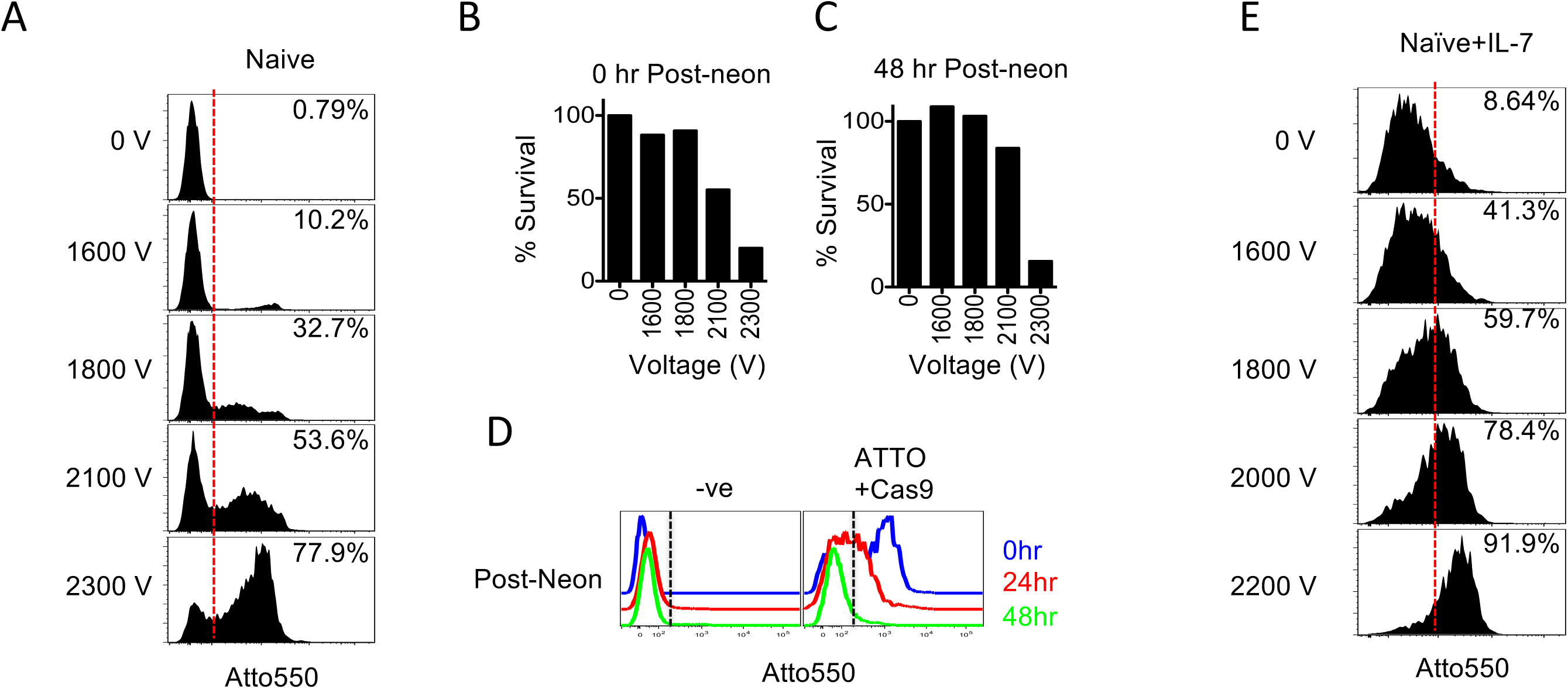
Naïve CD8 T cells need IL-7 stimulation to take up CAS9 complex following electroporation. A. Antigen-specific CD8 T cells were purified from naïve spleens and electroporated with ATTO550-tagged Cas9. Histograms depict uptake CAS9 complex by fluorescence of tracrRNA-ATTO550. B-C. Bar graph shows survival of antigen-specific CD8 T cells B. directly after electroporation or C. following electroporation and 48 hours of in vitro stimulation with APB-GP33+IL-2 for 48 hrs. D. Histograms depict amount of ATTO550 retained 0-48 hours post-electroporation. E. Antigen-specific CD8 T cells were purified from naïve spleens and cultured for 24hrs with IL-7. Cells were then electroporated with ATTO550-tagged Cas9 and indicated voltages. Percent in histogram represents amount of cells beyond the red dotted line.

**Supplementary Figure 3.**
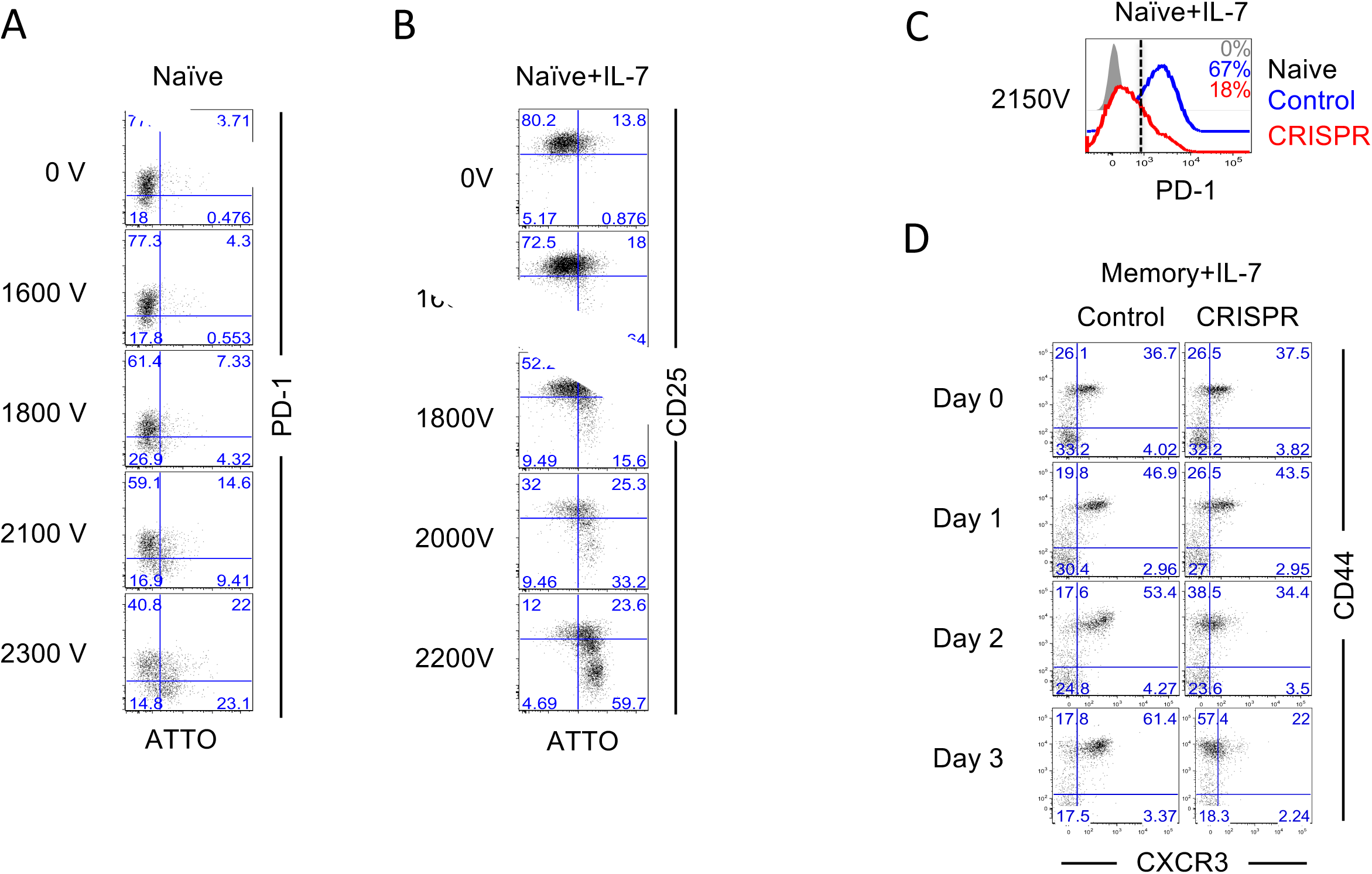
Incubation with IL-7 leads to optimal CRISPR mediated knockout in resting CD8 T cells. A. Naïve CD8 T cells were purified from spleens and electroporated with Cas9-PD1. Cells were then cultured in vitro with GP33-APB for 48 hours. B-C. Naïve CD8 T cells were purified from spleens and cultured for 24hrs with IL-7 in vitro. Cells were then electroporated with B. Cas9-CD25 or C. Cas9-PD-1. Cells were then stimulated in vitro with GP33-APB and target protein expression was measured 48hours following in vitro activation. D. Naïve B6 mice were infected with LCMV Arm. Spleens were collected from infected mice at day >60 post infection when memory has formed. CD8 T cells were purified from spleens and cultured for 24hrs in IL-7. Cells were then electroporated with Control or Cas9-CXCR3(CRISPR) at 2200V. Cells were transferred back in vitro and stimulated with IL-7 and IL-15. FACS plots depict CXCR3 expression as a function of CD44 at given times post neon transfection.

**Supplementary Figure 4.**
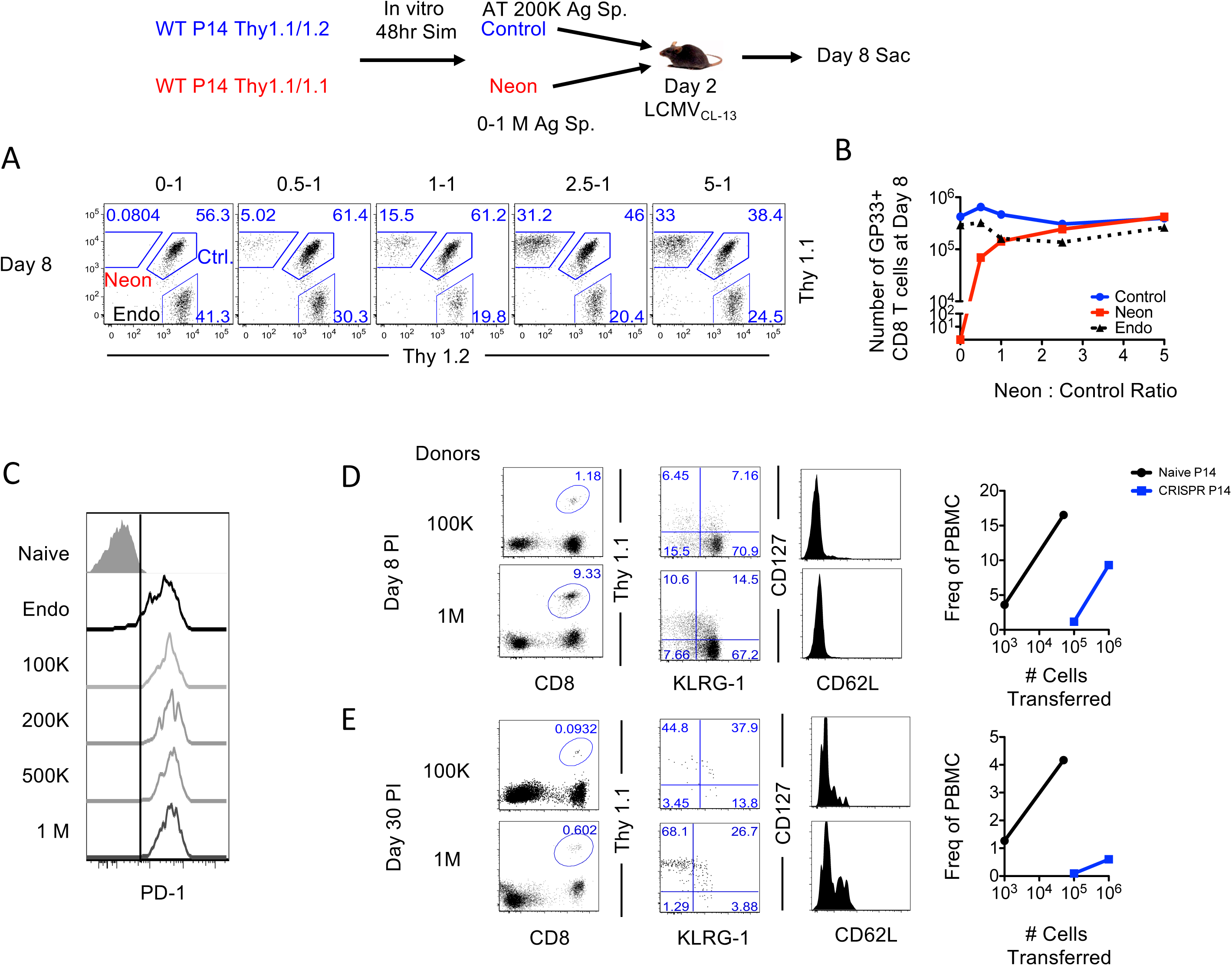
Titration of Donor CD8 T cells into infection matched mice to match endogenous CD8 T cell responses. Thy mismatched P14 CD8 T cells were purified from naïve mice and activated in vitro using GP33-APB for 48 hours. Cells were then electroporated using 1600V (Neon) or left unelectroporated (Control). Control and Neon cells were mixed together at given ratios and transferred into LCMVCL-13 infection matched mice. A. Mice were followed to peak of effector expansion at day 8 post-infection and spleens were collected. FACS plots are gated on GP33+CD44+ antigen-specific CD8 T cells. B. Line graph shows number of cells isolated from spleens of infected animals. C. Histograms are gated on Endogenous GP33 specific (Endo), or Neon Donor cells at day 8 post-infection. D-E. Antigen-specific CD8 T cells were activated for 48hrs in vitro and electroporated using the Neon instrument. Cells were immediately transferred into LCMV Arm infection matched mice at given amounts and followed to day 8 and 30 post infection. FACS plots depict frequency of donors in PBMC as well as the phenotype of donors. Line plot shows the relationship between transferred antigen-specific CD8 T cells and the observed frequency in PBMCs for Naïve cells transferred prior to LCMV infection (Black) or day 2 transferred CRISPR electroporated cells (Blue).

**Supplementary Figure 5.**
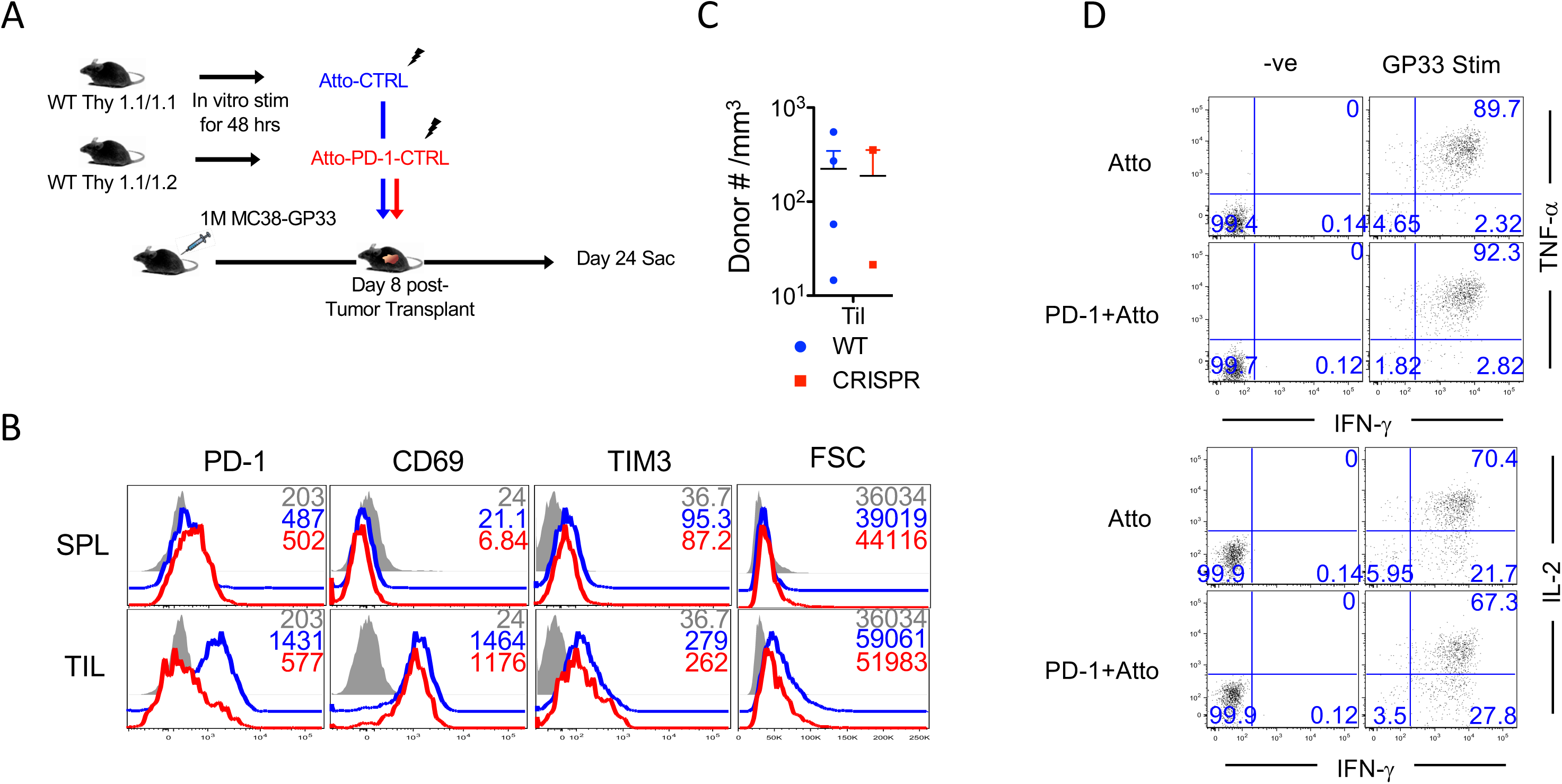
Distinct phenotypes of antigen-specific CD8 T cells in antigen free and tumor burdened environments. A. Naïve CD8 T cells were purified from spleens and stimulated in vitro for 48 hours using GP33-APB. Cells were electroporated with ATTO alone (control) or CAS9-PD-1-ATTO (PD-1 knockout) and transferred into tumor-burdened mice. B. Histograms are gated on donor antigen-specific CD8 T cells in spleen and TILs at day 24-post tumor transfer. Gray histogram is gated on endogenous CD44lo splenocytes. C. Graph shows number of donor CD8 T cells in TIL by volume of tumor. D. Splenocytes were stimulated for 5 hours in the presence of GP33 and BFA to measure cytokine functionality of donor cells. FACS plots are gated on donor CD8 T cells in spleen.

